# Mammal responses to habitat degradation induced by cashew expansion in West Africa

**DOI:** 10.1101/2025.03.29.646115

**Authors:** Daniel Na Mone, Isnaba Nhassé, João Soares, Raquel Oliveira, Manuel Lopes-Lima, Luís Palma, Ana Filipa Palmeirim

## Abstract

Tropical landscapes are perishing due to high rates of land-use change. In West Africa, Guinea-Bissau lost 77% of its closed-canopy forest over the last 25 years, mostly to the expansion of cashew monoculture. To date, understanding how species cope with such changes remains a conservation priority for the region. Here we examine mammal species composition, richness, and abundance, in addition to trophic-guild abundance across closed-canopy sub-humid forests and cashews orchards in the Cantanhez National Park, southwest Guinea-Bissau. Mammals were surveyed using one camera trap at each of the 24 sites, half in forest and half in cashew orchards, and their local and landscape-scale variables were measured. Based on 709 trap-nights, we collected 842 records from 25 mammal species. Local habitat structure—including canopy openness, floor obstruction, density of both lianas, palms and trees, tree species richness and height—sharply contrasted between forests and cashew orchards. As expected, mammal species composition differed between forests and cashew orchards, and the estimated, but not the observed, species richness declined towards cashew-like habitats. Although overall mammal abundance remained unaffected by the variables considered, carnivores were more abundant in cashew-like habitats, whereas insectivores and herbivores were more abundant in forest-like habitats. Human activity and distance to forest did not affect the response variables considered. Forest conversion into cashew orchards negatively affects mammals by profoundly changing the local habitat structure. Guild-level responses further unveiled specificity in their vulnerability to this form of land-use change, alerting for a potential disruption in the ecosystem functioning. We emphasize the need for policies that limit monoculture expansion, while safeguarding the remaining forests, thus maximising biodiversity persistence across the Afrotropics.

## Introduction

Tropical forests, holding over 75% of all species described to science, are under mounting pressure from agricultural expansion and intensification (IUCN, 2020; Banks-Leite *et al*., 2023). In West Africa, Guinea-Bissau is experiencing an unprecedent expansion of cashew cultivation (Catarino *et al*., 2015), having lost 77% of its closed-canopy forests between 2001 and 2018 (UN-FCCC, 2019). Today, the country’s economy strongly relies on the exportation of cashew nuts, which accounts for 90% of the GDP (UN-FCCC, 2019). Despite the potential pervasive impacts that this widespread land-use change might have on biodiversity (Guedes *et al*., 2024), evidence-based knowledge is largely unavailable across the Western Afrotropics (Luiselli *et al*., 2019).

The persistence of biodiversity in human-altered habitats is linked to the extent of transformation experienced by native habitats, which in turn determines the availability of food, structures for locomotion and resting, and changes in local climatic conditions (Gibson *et al*., 2011; Palmeirim *et al*., 2017). The higher the degree of habitat transformation, including its structure and composition, the more likely it is to be less suitable for the baseline species assemblages (Almeida-Maués *et al*., 2022). For example, the conversion of forests into cashew monocultures results in a significant loss of horizontal and vertical structural complexity, coupled with the periodic removal of understory vegetation (Temudo & Abrantes, 2014; Komanduri *et al*., 2023). Consequently, cashew orchards are expected to support a reduced number of species along with a markedly different species composition when compared to adjacent forests (Vasconcelos *et al*., 2015; Palmeirim *et al*., 2024). Additional factors affecting species persistence in human-modified landscapes may be related to landscape-scale characteristics, such as the proximity of source areas — i.e., habitats or regions that act as reservoirs that supply individuals to other areas via dispersal (Hansen & DeFries, 2007). Moreover, as wild animals are expected to avoid direct human contact (Happold, 1995), increased human activity could result in reduced wildlife habitat use (Rogala *et al*., 2011).

The complexity of species’ responses to land-use change depends on the species intrinsic characteristics (Newbold *et al*., 2019). For example, species within the same trophic guild often share similar ecological roles and resource requirements, leading them to interact with their environment in comparable ways and exhibit analogous responses to land-use change (Wearn *et al*., 2017). Trophic generalists such as omnivores may be particularly resilient to such changes due to their capacity to exploit a wider range of resources, enabling them to adapt more readily to altered environments (Law *et al*., 2017). Thus, more trophic-constrained species, such as carnivores, insectivores, and herbivores, may be generally more closely tied to habitat changes that affect the availability of prey and plants. Instead, trophic generalists, such as omnivores, might be more adaptable and less affected by such changes, due to their ability to rely on a wider range of resources (Law *et al*., 2017).

Tropical mammal assemblages are characterised by a remarkable taxonomic and functional diversity, thus playing critical roles in ecosystem processes. Mammals are involved in vital services that contribute to human well-being, including pollination, insect pest control, and bioturbation of soils (Lacher *et al*., 2019). As human land-use change intensifies, wild mammal populations typically decline (Ceballos *et al*., 2017) and species assemblages undergo significant shifts in composition (Foord *et al*., 2018). For example, the expansion of cashew plantations in India has led to the selective loss of native forest specialist mammals (Rege *et al*., 2020), while it has negatively affected habitat use by herbivores in northern Guinea-Bissau (Rossinyol-Fernàndez *et al*., 2024) and by several primate species in the southwest of the country (Bersacola *et al*., 2022).

Cantanhez National Park in south-western Guinea-Bissau contains one of the northernmost dense forests in West Africa, making it of high conservation importance (Dodman *et al*., 2004; Rainho *et al*., 2007). Despite being a protected area, 16% of its surface has already been occupied by cashew orchards (Pereira *et al*., 2022). However, the limited number of systematic studies carried out to date hinders our understanding of how this type of land-use change affects biodiversity (Guedes *et al*., 2024, but see Bersacola *et al*., 2022). To address this gap, we surveyed mammals at 24 sites, half of which located in forests and the other half in cashew orchards, in the Cantanhez National Park. We assessed mammal species composition, richness, and abundance (defined as the number of camera-trap records). In addition, we also analysed the abundance of carnivores, insectivores, omnivores, and herbivores. Given the expected contrasting local habitat structure between the forests and the cashew orchards (e.g., greater openness and lower height of the canopy, and lower tree species richness in the latter), we expected to observe different species composition between the two land-cover types, given the fewer mammal species and lower abundance in cashew-like habitats. Similarly, we expected mammal species richness and abundance to be lower in sites characterised by a greater distance from the closest forest and higher levels of human activity. We expected these patterns to be more pronounced when considering the abundance of carnivores, insectivores, or herbivores and less evident when considering omnivores.

## Methods

### Study area

The Cantanhez National Park (CNP) with an area of 1057 km^2^, was established in 2008 in the Tombali region of Guinea-Bissau (10°55’– 12°45’ N, 13°37’–16°43’ W; Fig. 1a). The CNP encompasses sub-humid closed-canopy forests, mangroves, wet grasslands, agricultural fields (mostly cashew orchards), and small human settlements (Pereira *et al*., 2022). The region has a humid monsoonal climate characterized by distinct dry (November to May) and wet (June to October) seasons. Average annual temperatures range from 28°C to 31°C, and annual rainfall reaches 2,400 mm (Climate Knowledge Portal, 2021).

**Figure 1.**
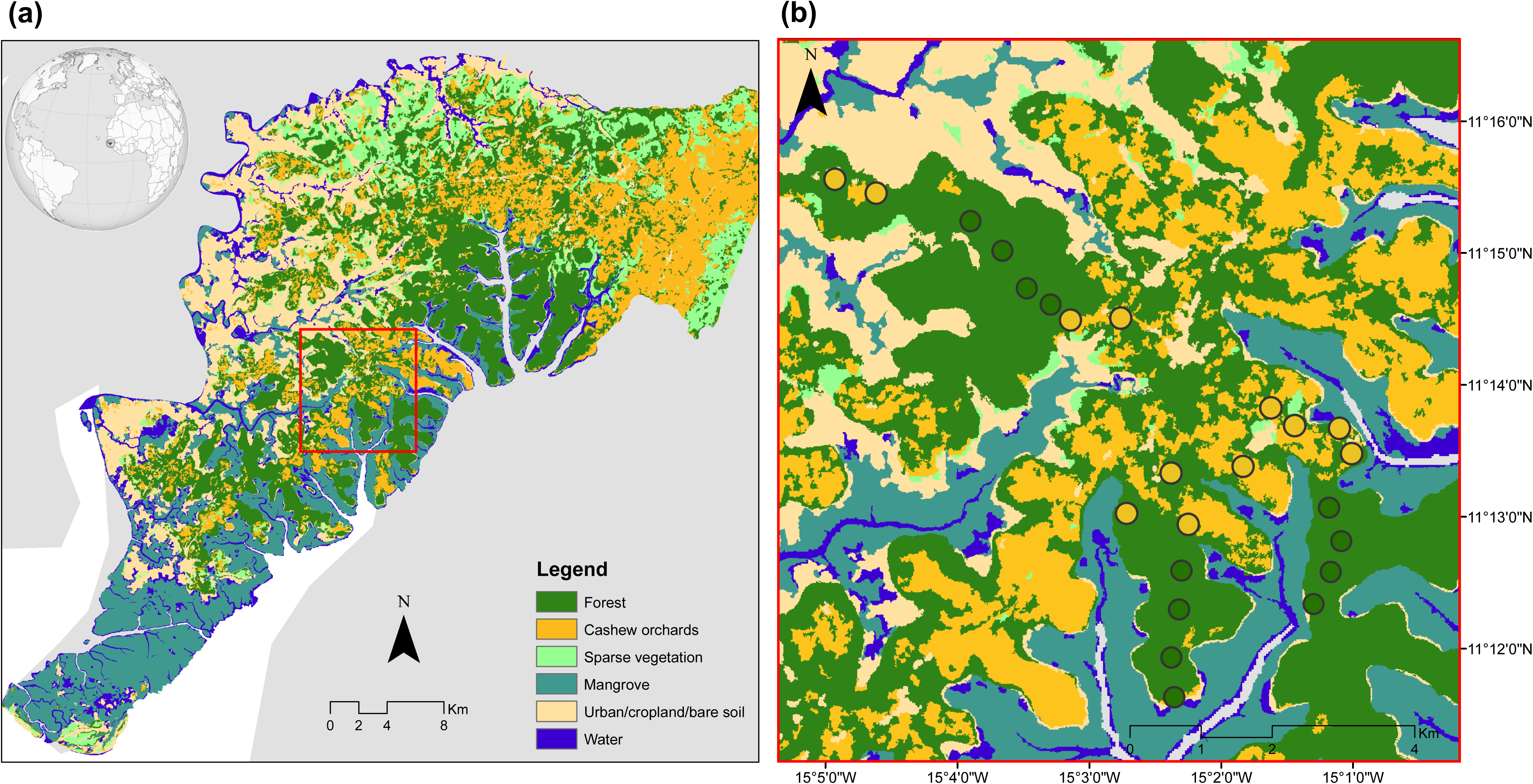
(a) Map of the Cantanhez National Park coloured according to the different types of land-use as extracted from Pereira et al. (2022) and location of the sampling sites within the study area (red square). (b) Location of the study area in detail, including the sampling sites located in the forest (green circles) and in the cashew orchards (orange circles).

In CNP, forests are often fragmented due to agricultural expansion and human settlements. However, the region retains patches of relatively intact forest, which play a vital role in maintaining wildlife corridors (Bersacola *et al*., 2022). Cashew orchards consist of monocultures of the Brazilian-native species *Anacardium occidentale* L., typically cultivated on small, locally managed plots. The trees, spaced 4 to 5 meters apart, are grown without the use of agrochemicals or irrigation. The cashew nut harvest takes place from early March to late June, while from October to December, the trees do not bear fruit, and the understory is characterized by low to moderate vegetation density. In December, farmers usually clear the orchards, removing undergrowth to prepare for the next season (Sierra-Baquero *et al*., 2024).

### Camera-trap surveys

Between November and December 2023, mammals were surveyed using camera-trapping in a central area of the CNP (Fig. 1a), where we established 24 sampling sites, half of which located in forests and the other half in cashew orchards. This area was characterised by a balanced forest-cashew land cover mosaic. The location of the sampling sites in the forests was related to the availability of large forest remnants in that part of the park, whereas the location of the sites in the cashew orchards aimed to be in the vicinities of the sampling sites in the forests (Fig. 1b). This period, corresponding to the beginning of the dry season, was chosen due to the generally higher mammal activity during that part of the year (Rossinyol-Fernàndez *et al*., 2024). In each sampling site, we deployed one digital camera (Browning Patriot model BTC-PATRIOT-FHD) that was continuously operating for 30 days. Within each land cover type, the location where each camera-trap was deployed was carefully selected by an experienced local nature guide (Mamadu Cassamá) aiming to maximise the detection of mammal species, thereby ensuring a reasonable visibility in the camera detection range. Whenever possible, camera-trap location also avoided proximity to trails used by people. We configured each camera to obtain a sequence of five photographs with a 15-second interval. Deployed cameras were unbaited, attached to a tree trunk, 30 – 40 cm above ground, and spaced at least 300 m from adjacent sampling sites. The camera’s detection range (approximately 5 × 5 m in front of the camera) was cleared by the time the cameras were deployed.

The collected photographs were processed using TimeLapse Software (Greenberg *et al*., 2019; Greenberg, 2023), and each image was carefully reviewed to identify mammals to the species level, whenever possible. Consecutive photographs of the same species taken within a 30-minute interval were considered a single detection event (hereafter referred as a species *record*) (Gessner *et al*., 2014) unless individuals could be distinguished by clear differences in age, sex, or unique morphological traits. Due to poor photo quality or partial capture of the individual, 16 out of 858 records (1.9%) could not be positively identified and were excluded from further analysis. We considered the number of records as a proxy of species abundance (e.g., Rossinyol-Fernàndez *et al*., 2024 and Goebel *et al*., 2025). Given logistical difficulties that caused a slightly different number (between 15 and 30) of trapping days between sites (Table S1), we used the standardised species abundance given by the number of records per 10 trap-days (hereafter, species abundance). The overall sampling effort across sites amounted to 709 camera-trap days (Table S1). Fieldwork was conducted under permission from the national biodiversity and protected areas authority, the ‘Instituto da Biodiversidade e das Áreas Protegidas’ (IBAP). In addition, permission was obtained from the owners of all cashew farms included in the survey prior to sampling.

### Local habitat structure, distance to forest and human activity

To explain mammal responses to cashew expansion, we considered variables related to the local habitat structure, landscape, and degree of human activity. As for the local habitat structure, we measured canopy openness, floor and understorey obstruction, density of lianas, palms, and hardwood trees with diameter at breast height (DBH) above 10 cm, tree species richness and height. Variables requiring visual estimates were consistently undertaken by the same observer, whereas tree species identification was done by the local nature guide (Mamadu Cassamá). A Principal Component Analysis (PCA) was applied to illustrate how the local habitat variables vary between land cover types, using the ‘ggbiplot’ R package (Vu *et al*., 2011). The scores of the PC1—synthesising the local habitat structure of each sampling site—were retained, comprising the local habitat structure variable included in subsequent models. Distance to forest refers to the closest Euclidean distance between the camera-trap and the edge of the closest forest and was measured using Google Earth imagery (Google Earth v7.3.6.9345, 2022). The degree of human activity was given by the number of camera-traps records of human presence in the sampling sites. These photos were first separated from those containing non-human records, a blur filter was applied to exclude the recognition of humans, and the number of photos was counted. Consecutive photographs taken within a 30-minute interval were considered a single detection event. The number of records was standardised per 10 trap-days. All variables are described in Table 1.

**Table 1.**
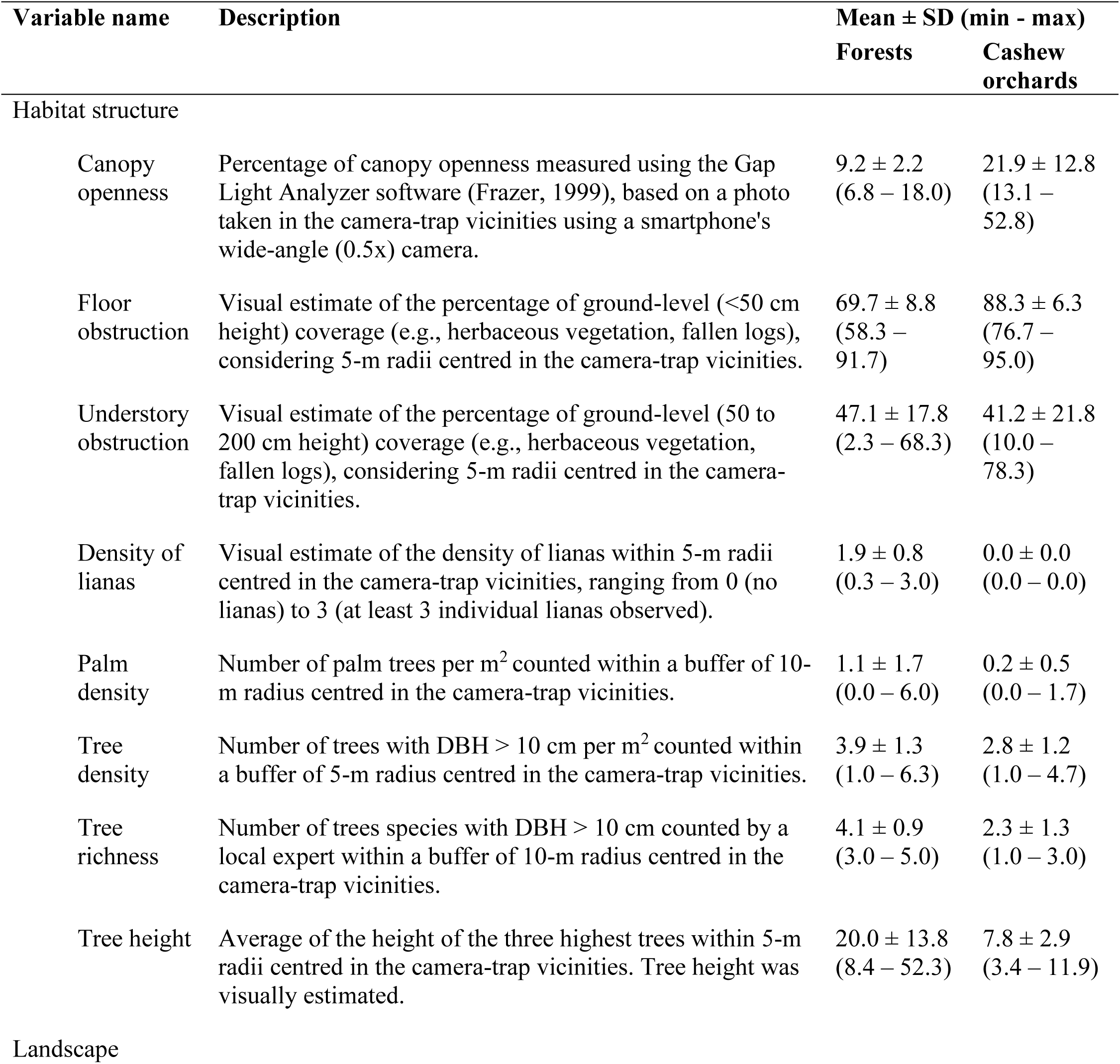
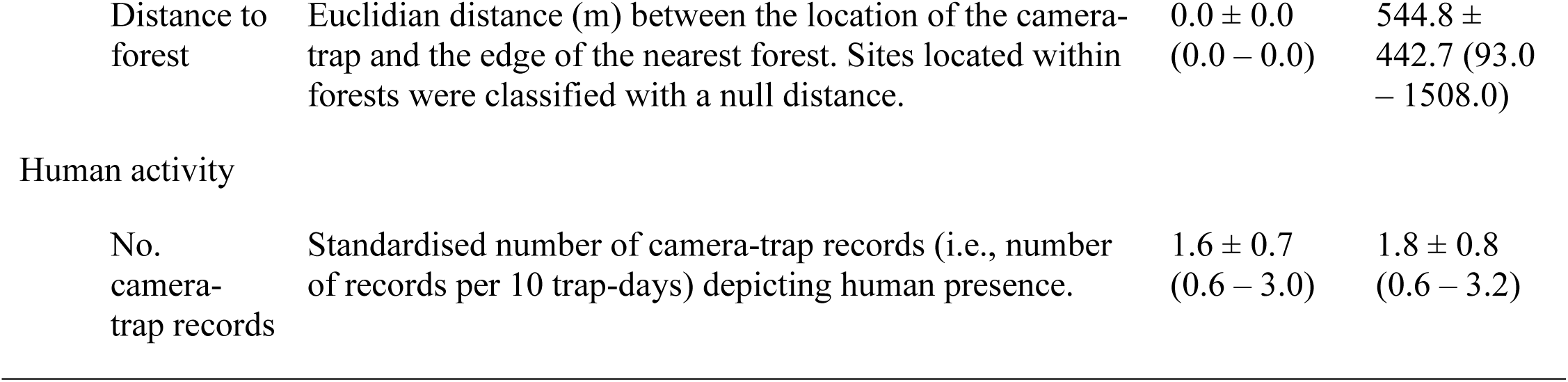
Description of the habitat structure, landscape and human activity variables used to predict mammal diversity in the Cantanhez National Park, Guinea-Bissau. The average, standard deviation, minimum and maximum values are provided for sampling sites located within forests and within cashew orchards.

| Variable name | Description | Mean ± SD (min - max) | |
| --- | --- | --- | --- |
|  |  | Forests | Cashew orchards |
| Habitat structure |  |  |  |
| Canopy openness | Percentage of canopy openness measured using the Gap Light Analyzer software (Frazer, 1999), based on a photo taken in the camera-trap vicinities using a smartphone's wide-angle (0.5x) camera. | 9.2 ± 2.2<br>(6.8 – 18.0) | 21.9 ± 12.8<br>(13.1 – 52.8) |
| Floor obstruction | Visual estimate of the percentage of ground-level (<50 cm height) coverage (e.g., herbaceous vegetation, fallen logs), considering 5-m radii centred in the camera-trap vicinities. | 69.7 ± 8.8<br>(58.3 – 91.7) | 88.3 ± 6.3<br>(76.7 – 95.0) |
| Understory obstruction | Visual estimate of the percentage of ground-level (50 to 200 cm height) coverage (e.g., herbaceous vegetation, fallen logs), considering 5-m radii centred in the camera-trap vicinities. | 47.1 ± 17.8<br>(2.3 – 68.3) | 41.2 ± 21.8<br>(10.0 – 78.3) |
| Density of lianas | Visual estimate of the density of lianas within 5-m radii centred in the camera-trap vicinities, ranging from 0 (no lianas) to 3 (at least 3 individual lianas observed). | 1.9 ± 0.8<br>(0.3 – 3.0) | 0.0 ± 0.0<br>(0.0 – 0.0) |
| Palm density | Number of palm trees per m<sup>2</sup> counted within a buffer of 10-m radius centred in the camera-trap vicinities. | 1.1 ± 1.7<br>(0.0 – 6.0) | 0.2 ± 0.5<br>(0.0 – 1.7) |
| Tree density | Number of trees with DBH > 10 cm per m<sup>2</sup> counted within a buffer of 5-m radius centred in the camera-trap vicinities. | 3.9 ± 1.3<br>(1.0 – 6.3) | 2.8 ± 1.2<br>(1.0 – 4.7) |
| Tree richness | Number of trees species with DBH > 10 cm counted by a local expert within a buffer of 10-m radius centred in the camera-trap vicinities. | 4.1 ± 0.9<br>(3.0 – 5.0) | 2.3 ± 1.3<br>(1.0 – 3.0) |
| Tree height | Average of the height of the three highest trees within 5-m radii centred in the camera-trap vicinities. Tree height was visually estimated. | 20.0 ± 13.8<br>(8.4 – 52.3) | 7.8 ± 2.9<br>(3.4 – 11.9) |
| Landscape |  |  |  |
| Distance to forest | Euclidian distance (m) between the location of the camera-trap and the edge of the nearest forest. Sites located within forests were classified with a null distance. | $0.0 \pm 0.0$<br>(0.0 – 0.0) | $544.8 \pm 442.7$ (93.0 – 1508.0) |
| Human activity |  |  |  |
| No. camera-trap records | Standardised number of camera-trap records (i.e., number of records per 10 trap-days) depicting human presence. | $1.6 \pm 0.7$<br>(0.6 – 3.0) | $1.8 \pm 0.8$<br>(0.6 – 3.2) |

### Data analysis

We assessed sampling sufficiency for each site by examining rarefaction curves using the ‘iNEXT’ R package (Hsieh *et al*., 2016). Rarefaction curves, denoting sampling sufficiency at the site level, did not stabilize in any of the sampling sites (Fig. S1). Following Chao et al. (2014), to account for the incomplete sampling sufficiency of some sites (e.g., cam3 and cam4), we estimate species richness by applying the richness estimator available in the same R package, which accounts for twice the number of records in each site. Two sites were surveyed with a particularly low sampling effort, namely “cashew3” (24 trap-nights) and “cashew7” (15 trap-nights). In these sites, only four and two records each, of one and two species, respectively, were obtained. As the exceptional low number of records in these sites might be due to the low sampling effort, we considered those sites as outliers, and those were removed from subsequent analyses.

Species composition across the different land cover types was examined using a Non-Metric Multidimensional Scaling (NMDS) ordination based on a Bray-Curtis similarity matrix, considering the number of records per site for each species (stress = 0.149). We tested whether species composition varied between land cover types using a Permutational Multivariate Analysis of Variance (PERMANOVA). Complementarily, we applied a Permutational Analysis of Multivariate Dispersions (PERMDISP) to compare the dispersion in species composition between land cover types. These analyses were carried out using the vegan R package (Oksanen *et al*., 2013).

We then analysed the combined effects of local habitat structure, distance to forest and degree of human activity on both observed and estimated species richness, overall species abundance and each trophic guild-level abundance, namely carnivores, insectivores, omnivores, and herbivores. Each species was classified according to either of these four trophic guilds based on their diet as extracted from the PanTHERIA (Jones *et al*., 2009) and Elton databases (Wilman *et al*., 2014) (Table S2). We performed Generalised Linear Models (GLMs) for each of these response variables (i.e., observed and estimated species richness, overall species abundance and abundance of carnivores, insectivores, omnivores, and herbivores). An offset variable regarding the log_10_ of the number of records (i.e., overall species abundance) was also included in the model regarding the estimated species richness, to account for the fact that a higher species richness might be due to the higher number of records. The decimal numbers in all response variables—except the observed species richness—were rounded to obtain a discrete value. All models were fitted with a Negative Binomial distribution. We tested for multicollinearity by calculating the Variance Inflation Factor (VIF) of each independent variable. No variables were found to be redundant (VIF > 4) (Dormann *et al*., 2013). A candidate model set was further constructed using all additive combinations of the three explanatory variables retained, and models were ranked based on their Akaike Information Criteria corrected for small sample size (AICc), using the ‘MuMIn’ R package (Bartoń, 2016). Whenever no model was performing as the ‘best’ model (i.e., ΔAICc < 2, with ΔAIC = AIC*_i_* − AIC_min_ in which *i* = *i*^th^ model), we performed a model-averaging approach using the entire set of candidate models to account for model uncertainty in multi-model inference. Explanatory variables were previously normalised to place coefficient estimates onto the same scale. We examined the residuals of each model using the ‘DHARMa’ R package (Hartig, 2022) and further applied a Moran I test to the residuals of each model to assess spatial autocorrelation using ‘spdep’ R package (Bivand *et al.,* 2017). Spatial autocorrelation was not detected in the residuals of any model. All analyses were conducted in R version 4.1.2 (R Core Team, 2021).

## Results

The characteristics of the local habitat structure sharply contrasted between forests and cashew orchards (Fig. 2). According to the PCA diagram, sampling sites located in forests were characterised by higher values of tree height, liana density, tree density and tree species richness, whereas cashew orchards were mostly characterised by higher values of floor obstruction, canopy openness and palm density.

**Figure 2.**
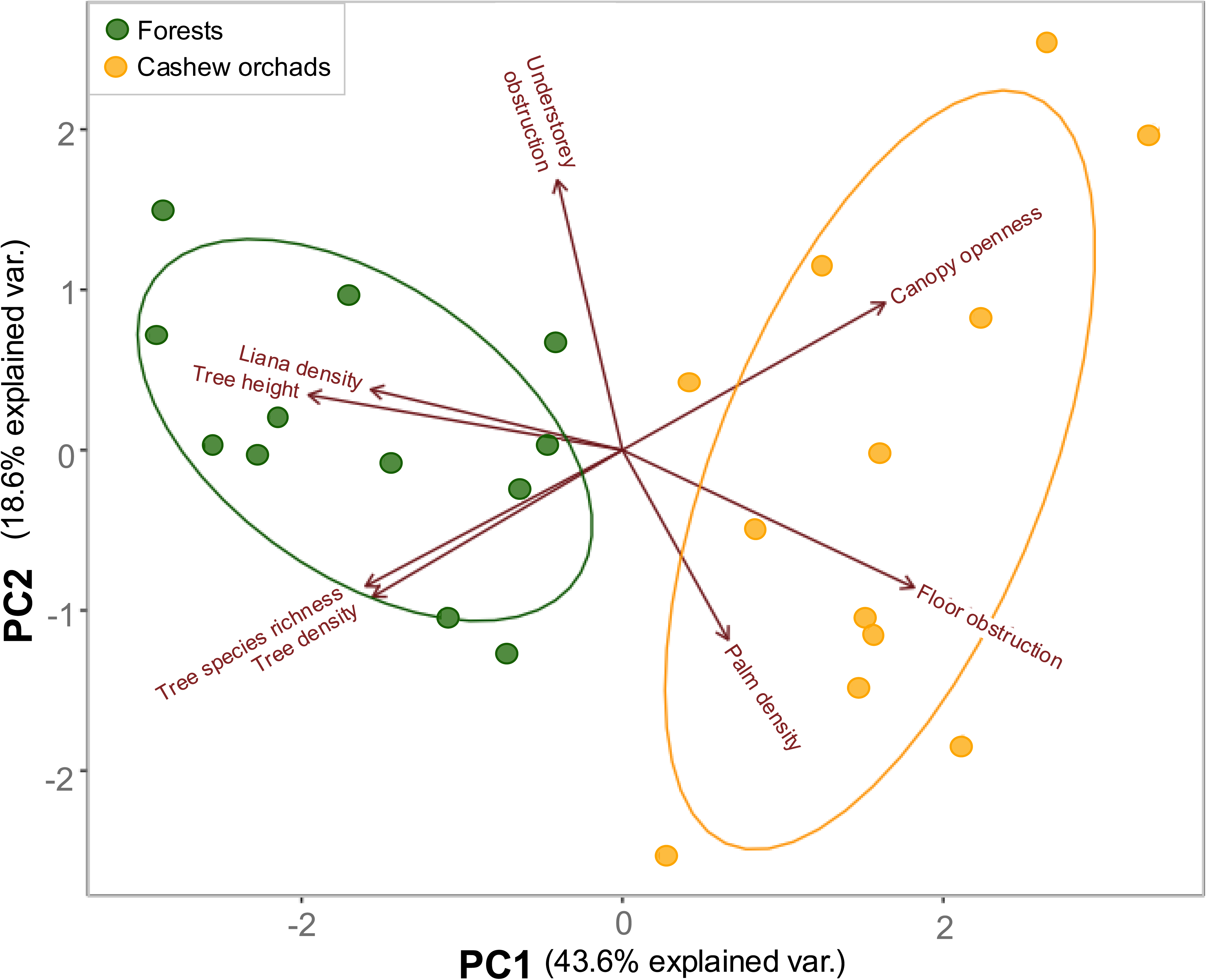
Diagram of the Principal Component Analysis (PCA) based on the characteristics of each of the 24 sampling sites located in closed-canopy forests and cashew orchards in the Cantanhez National Park, Guinea-Bissau. Sampling sites are colour-coded according to the land-use type where the site was located. The strength of each habitat characteristic considered is represented by the red arrows and the name of the habitat characteristic is provided (a description of each habitat characteristic can be found in Table 1). The variance explained by each axis (i.e., PC1 and PC2) is correspondingly indicated in the plot.

In total, we obtained 842 records of 25 mammal species, corresponding to 12 families and four orders. These species included seven carnivore (*n* = 171 records, 20.3%), four insectivore (*n* = 219, 26.0%), nine omnivore (n = 403, 47.9%) and five herbivore species (n = 49, 5.8%) (Table S2). The most recorded species was the giant pouched-rat *Cricetomys gambianus* (29.7%), followed by the white-tailed mongoose *Ichneumia albicauda* (23.3%). Seven species, namely the marsh cane-rat *Thryonomys swinderianus*, the red river hog *Potamochoerus porcu*s, the banded mongoose *Mungos mungo*, the yellow-backed duiker *Cephalophus silvicultor*, the Gambian sun squirrel *Heliosciurus gambianus*, the Egyptian mongoose *Herpestes ichneumon*, and the honey badger *Mellivora capensis*, were recorded only once or twice throughout the sampling (Table S2). An average (± SD) of 8.1 ± 2.6 species was recorded in forests (60.7% of the records), and 6.1 ± 3.1 species in cashew orchards (39.3%).

Six species were exclusively recorded in forests and another six species only in cashew orchards (Fig. 3a). From the 14 species using both land cover types, eight species had a higher percentage of records within forests, while four had higher percentage of records within cashew orchards (Fig. 3a). Despite some overlap observed in the NMDS diagram (Fig. 3b), species composition differed between forests and cashew orchards (*P* = 0.033, Table S3). However, no differences were observed in the dispersion of species composition (Table S3).

**Figure 3.**
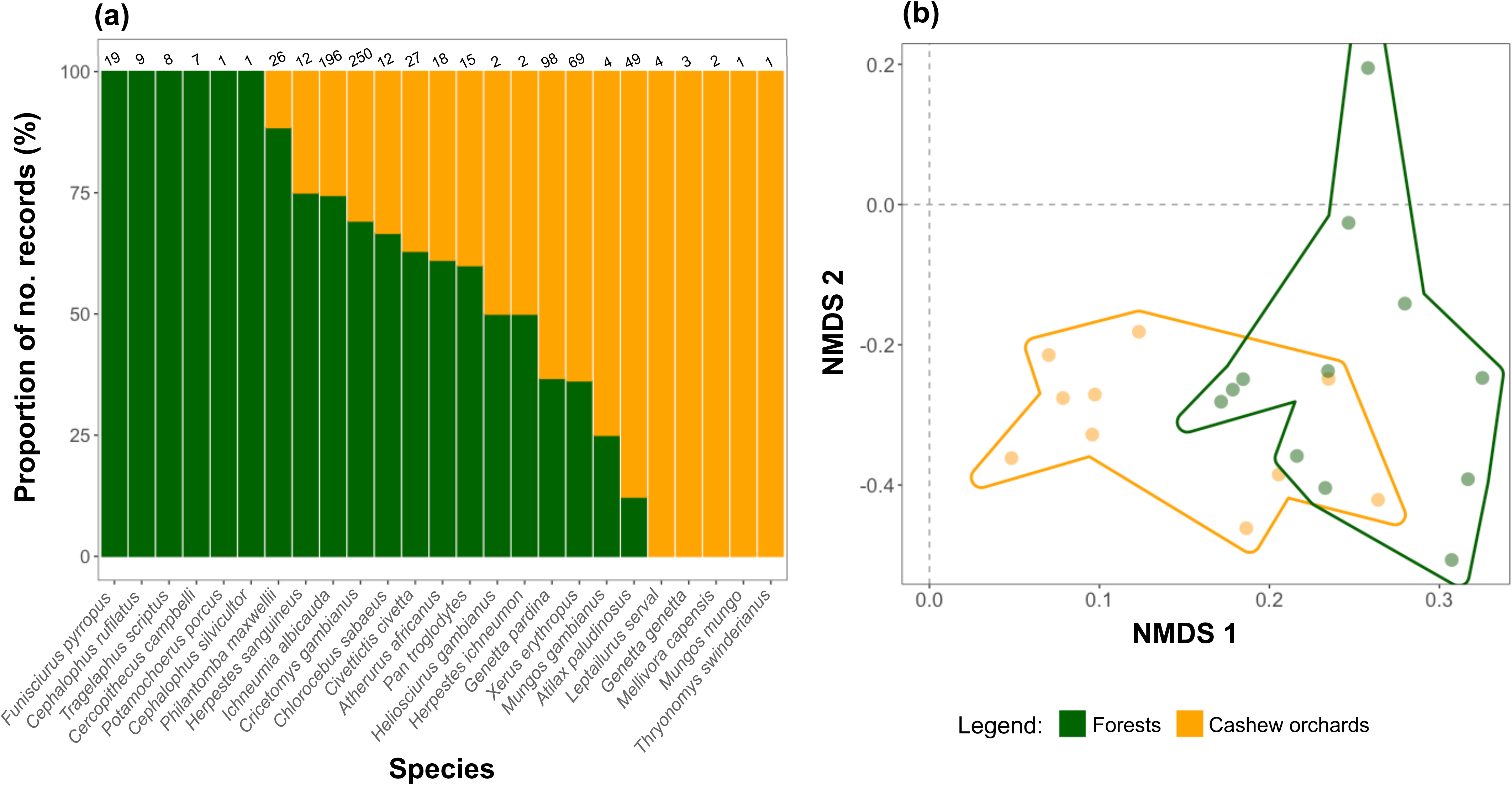
(a) Proportion of mammal species records obtained in forest patches (in green) and in cashew orchards (in orange); the numbers in brackets correspond to the total number of records of each species; species are displayed from highest to lowest proportion of records obtained in forests; (b) non-metric dimensional scaling (NMDS) representing mammal species composition across forest patches (in green) and cashew orchards (in orange); this corresponded to 24 sampling sites in the Cantanhez National Park, Guinea-Bissau.

The top-ranked models explaining the observed species richness included the null model (AICc = 107.0) and the one including only human activity (AICc = 108.3); those explaining the estimated species richness included either only habitat structure or both habitat structure and human activity (AICc = 134.7 and 135.0, respectively). As for the overall species abundance, the null model performed the best (AICc = 149.2). When considering separately each of the trophic guilds, carnivore abundance was best explained by the model including only the habitat structure (AICc = 92.1); the top-ranked models for insectivore abundance included both distance to forest and habitat structure and only habitat structure (AICc = 102.5 and 102.7, respectively); those for omnivores included both the null model (AICc = 123.7) and the one including only human activity (AICc = 124.6); and those for herbivores included both the habitat structure (AICc = 48.8) and the null model (AICc = 50.8; Table S4).

Based on the model averaging/‘best’ model results (i.e., ΔAICc < 2), the observed species richness was not predicted by any of the variables considered (Table 2, Fig. S3). Yet, estimated species richness was affected by the habitat structure, being associated with negative scores of the PC1 (*β*_habitat_ = –0.191, *P* = 0.051, Table 2, Fig. 4a), which in turn is evidence of forest-like habitat characteristics. Overall species abundance was not predicted by any of the variables considered (Table 2, Fig. 4b). Yet, when considered at the trophic guild-level, the abundance of carnivores increased in habitats characterised by higher PC1 scores (*β*_habitat_ = 0.590, *P* = 0.004, Table 2, Fig. 4c), denoting characteristics coincident with cashew orchards. Omnivore abundance remained unaffected by the variables considered, whereas the abundance of both insectivores and herbivores decreased with the PC1 scores, i.e., in the direction of the cashew-like habitat characteristics (insectivores: *β*_habitat_ = –0.704, *P* = 0.016, Fig. 4e; herbivores: *β*_habitat_ = –0.779, *P* = 0.055, Fig. 4f).

**Figure 4.**
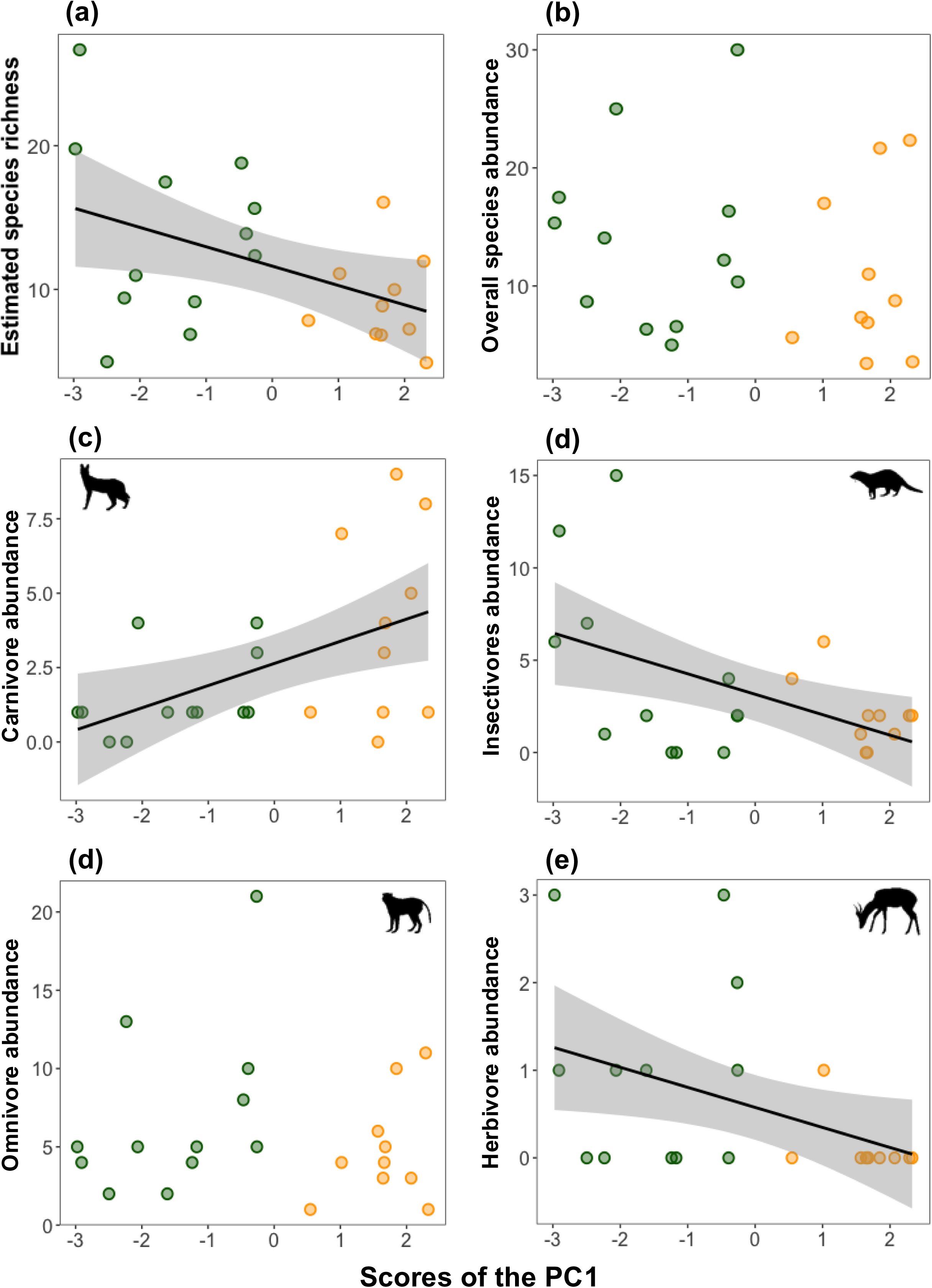
Relationship between (a) estimated species richness, (b) overall species abundance, (c) carnivore abundance, (d) insectivore abundance, (e) omnivore abundance and (f) herbivore abundance and the local habitat characteristics as denoted by the scores of the first component of the Principal Component Analysis (CP1). Shaded areas represent the 95% confidence region; solid dots indicate observed values; lines represent the model adjusted for the strongest relationships (*P* ≤ 0.05). Dots are colour coded according to the land-cover type where the sampling sites were located (forests in green and cashew orchards in orange).

**Table 2.**
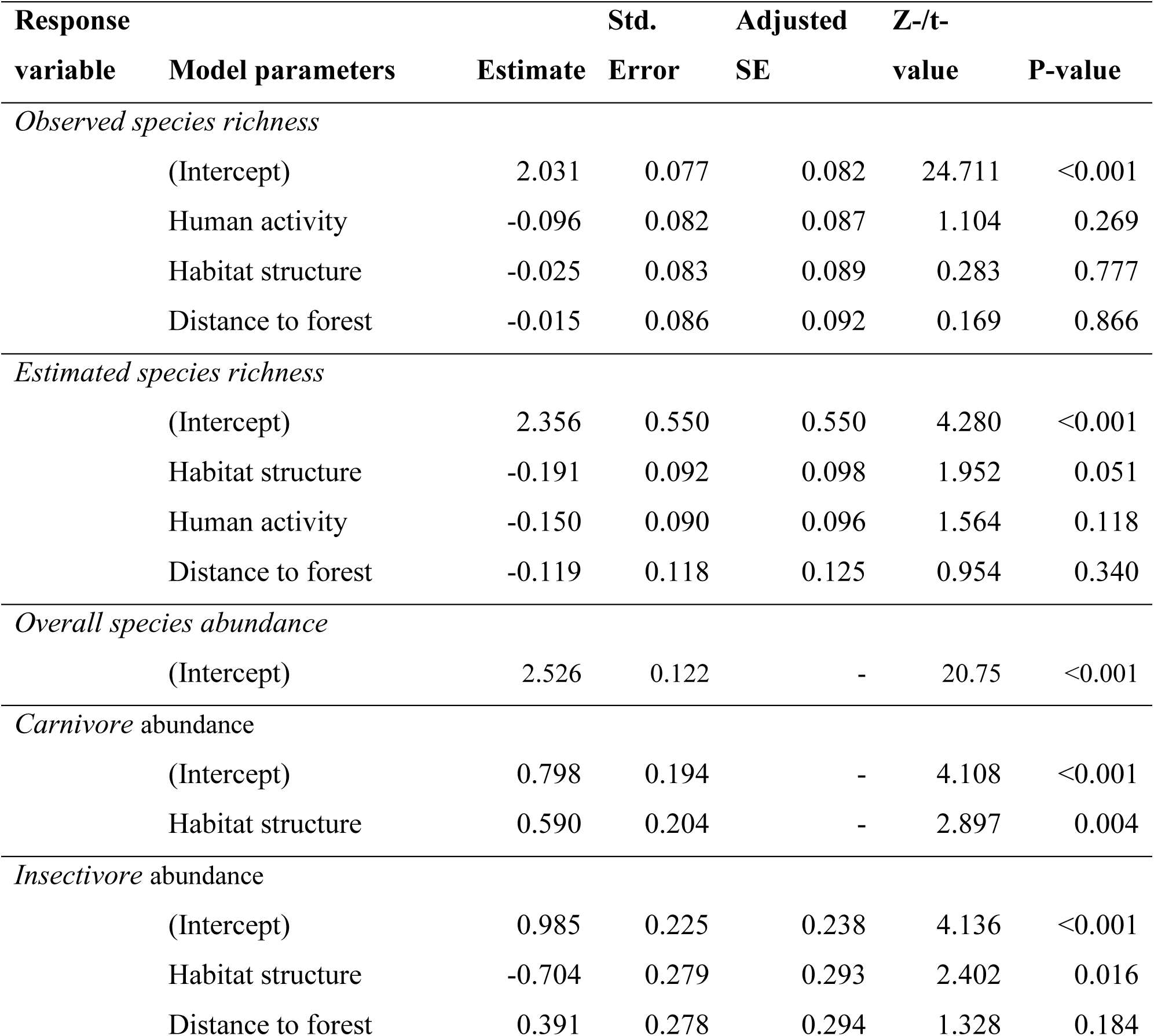
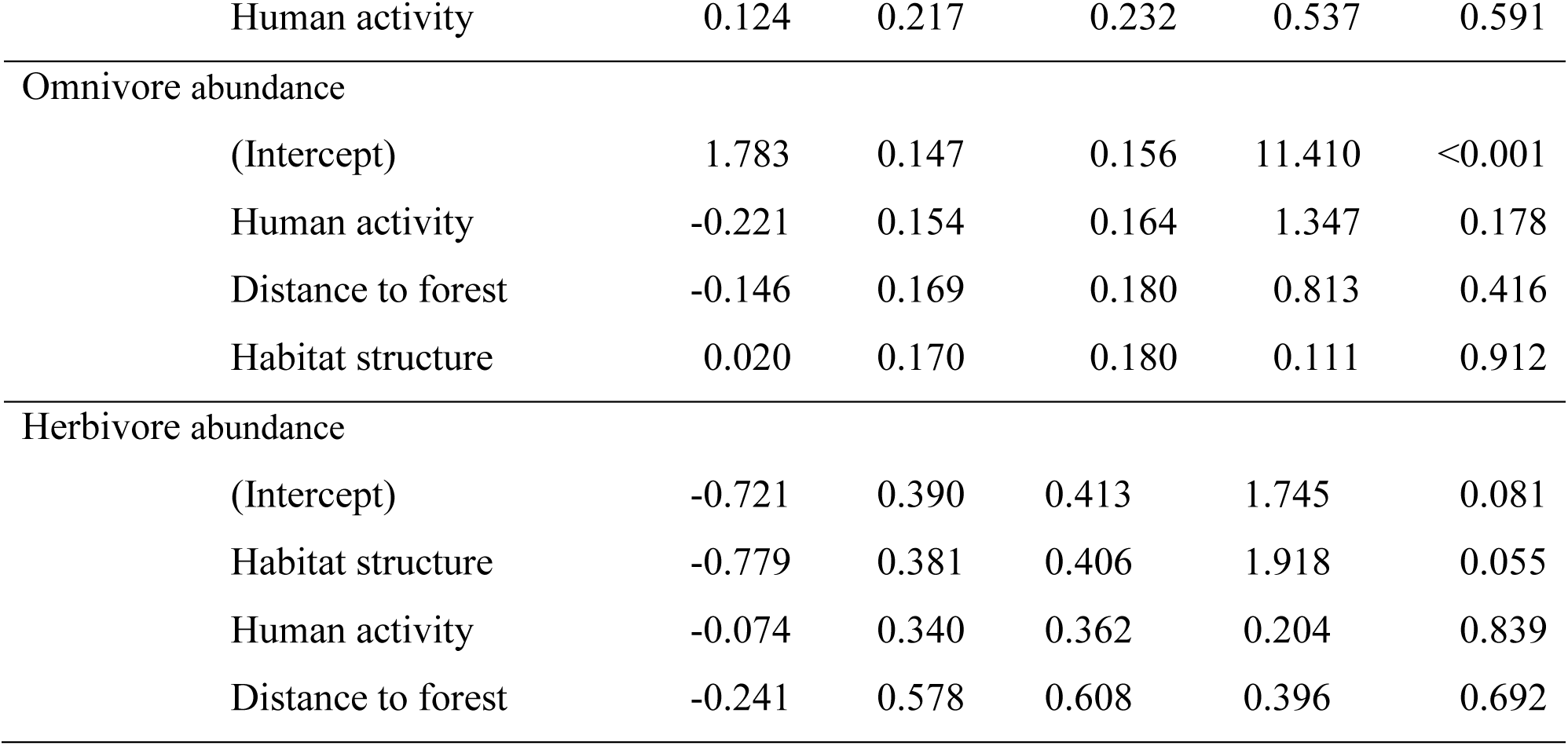
Summary of the averaged Generalized Linear Models (GLMs) relating observed and estimated mammal species richness, overall species abundance and abundance of each trophic guild—carnivorous, insectivorous, omnivorous, and herbivorous—according to habitat structure (Principal Component 1 of the PCA including the variables described in Table 1), landscape (distance to the nearest forest) and degree of human activity (standardised number of camera-trap records depicting human activity) across 24 sampling sites in the Cantanhez National Park, Guinea-Bissau. For each model, we indicate the estimate, standard error, adjusted standard error, Z-/t-values and P-value. Models were averaged considering the entire set of alternative models. As an exception, the models regarding overall species abundance and carnivore abundance corresponds to the best model since only one model showed ΔAICc < 2. For this model, t-value, instead of a z-value, is provided. The full set of alternative models and respective parameters can be found in Table S2.

| **Response variable** | **Model parameters** | **Estimate** | **Std. Error** | **Adjusted SE** | **Z-/t-value** | **P-value** |
| --- | --- | --- | --- | --- | --- | --- |
| *Observed species richness* |  |  |  |  |  |  |
|  | (Intercept) | 2.031 | 0.077 | 0.082 | 24.711 | <0.001 |
|  | Human activity | -0.096 | 0.082 | 0.087 | 1.104 | 0.269 |
|  | Habitat structure | -0.025 | 0.083 | 0.089 | 0.283 | 0.777 |
|  | Distance to forest | -0.015 | 0.086 | 0.092 | 0.169 | 0.866 |
| *Estimated species richness* |  |  |  |  |  |  |
|  | (Intercept) | 2.356 | 0.550 | 0.550 | 4.280 | <0.001 |
|  | Habitat structure | -0.191 | 0.092 | 0.098 | 1.952 | 0.051 |
|  | Human activity | -0.150 | 0.090 | 0.096 | 1.564 | 0.118 |
|  | Distance to forest | -0.119 | 0.118 | 0.125 | 0.954 | 0.340 |
| *Overall species abundance* |  |  |  |  |  |  |
|  | (Intercept) | 2.526 | 0.122 | - | 20.75 | <0.001 |
| *Carnivore abundance* |  |  |  |  |  |  |
|  | (Intercept) | 0.798 | 0.194 | - | 4.108 | <0.001 |
|  | Habitat structure | 0.590 | 0.204 | - | 2.897 | 0.004 |
| *Insectivore abundance* |  |  |  |  |  |  |
|  | (Intercept) | 0.985 | 0.225 | 0.238 | 4.136 | <0.001 |
|  | Habitat structure | -0.704 | 0.279 | 0.293 | 2.402 | 0.016 |
|  | Distance to forest | 0.391 | 0.278 | 0.294 | 1.328 | 0.184 |

|  |  |  |  |  |  |
| --- | --- | --- | --- | --- | --- |
| Human activity | 0.124 | 0.217 | 0.232 | 0.537 | 0.591 |
| Omnivore abundance |  |  |  |  |  |
| (Intercept) | 1.783 | 0.147 | 0.156 | 11.410 | <0.001 |
| Human activity | -0.221 | 0.154 | 0.164 | 1.347 | 0.178 |
| Distance to forest | -0.146 | 0.169 | 0.180 | 0.813 | 0.416 |
| Habitat structure | 0.020 | 0.170 | 0.180 | 0.111 | 0.912 |
| Herbivore abundance |  |  |  |  |  |
| (Intercept) | -0.721 | 0.390 | 0.413 | 1.745 | 0.081 |
| Habitat structure | -0.779 | 0.381 | 0.406 | 1.918 | 0.055 |
| Human activity | -0.074 | 0.340 | 0.362 | 0.204 | 0.839 |
| Distance to forest | -0.241 | 0.578 | 0.608 | 0.396 | 0.692 |

## Discussion

While tropical forests continue to face significant threats from agricultural expansion (Hansen *et al*., 2020), understanding how biodiversity responds to these land-use changes is essential for informing evidence-based conservation strategies (Caro *et al*., 2022). In this study, we highlight the widespread impacts of monoculture expansion in this under-studied region of West Africa. Our study found that cashew orchards, by substantially altering local habitat structure, support distinct mammal assemblages characterised by a lower species richness compared to nearby forests. As expected, insectivorous and herbivorous mammals were less often detected in cashew orchards. Notwithstanding, carnivores were more often detected in the cashew orchards, while omnivores were similarly recorded in forests and cashew orchards.

Undisturbed tropical forests are characterised by a complex vertical stratification, including towering emergent trees (Kricher, 2011). In contrast, human-modified land cover types exhibit significant compositional and structural differences that usually result in altered resources (e.g., food and shelter) and microclimatic conditions, such as increased temperatures and decreased humidity (Hardwick *et al*., 2015). These changes often result in reduced resource availability for forest-dependent species, contributing to the lower mammal species richness observed in the habitat-simplified cashew orchards. For example, the reduction in vertical complexity caused by forest conversion into cashew plantations— including a lower density of trees and lianas—has affected scansorial species such as the fire-footed rope squirrel *Funisciurus pyrropus* and the Campbell’s monkey *Cercopithecus campbelli*, both of which were absent from cashew orchards but were commonly found in nearby forests (but see Bersacola *et al*. (2022) for a detailed analysis of primate occupancy patterns in the CNP). Additionally, the increased canopy openness in cashew orchards has encouraged the growth of palm trees and herbaceous vegetation at the floor level. Even the most frequently recorded species, such as the herpestid *I. albicauda* and the rodent *C. gambianus*, predominantly used forest habitats, being observed in forests in approximately 75% of instances. This highlights the importance of maintaining forest patches even within modified landscapes to support biodiversity.

Guinea-Bissau is located within the Guinea-Congolese/Sudanese regional transition zone, where the dominant vegetation types include open forest and secondary formations such as woodland savannah, shaped by fire and shifting agriculture (Swaine, 1992), but with patches of dense forest formations that closely resemble the dense Guinean rainforests (Catarino & Indjai, 2019), as in the CNP area. We found a great structural disparity between forests and cashew orchards, leading to increased conservation value of these forests for certain species. In other systems, the structural contrast between forests and cultivated areas is less pronounced. This is the case of the northern region of Guinea-Bissau, characterised by a mosaic landscape of open forest patches traditionally managed by local communities (Palmeirim *et al*., 2023), along with rice fields and cashew orchards. Here, only minor, or negligible effects of forest conversion into cashew orchards have been reported for medium-sized mammals (Rossinyol-Fernàndez *et al*., 2024), small mammals (Oliveira *et al*., 2024), and herpetofauna (dos Reis-Silva *et al*., 2025). Similarly, Amazonian mammal assemblages are more severely impacted by forest conversion into structurally contrasting tree monocultures, while selectively logged and secondary regrowth forests support assemblages of higher ecological integrity (Almeida-Maués *et al*., 2022). Studies from India also highlight that both mammals and anurans frequently use cashew orchards (Rege *et al*., 2020), reinforcing that biodiversity responses to monoculture expansion may vary geographically.

Despite the lower estimated number of species using cashew orchards, more than half of the species were able to make use of cashew orchards to some extent. In this regard, we highlight that cashew orchards in Guinea-Bissau are not subject to chemical treatment and remain minimally managed (Monteiro *et al*., 2017; Sierra-Baquero *et al*., 2024). Indeed, human activity was low within cashew orchards (1.8 ± 0.8 records), at least during the sampling period (i.e., not overlapping the cashew fruiting season), and also in forests (1.7 ± 0.7 records). If organic and poor management of cashew orchards in Guinea-Bissau may not be beneficial for cashew productivity, it may favour their use by mammals.

In addition, five species were recorded exclusively in the cashew orchards, while another four species used cashew orchards more frequently than forests. Some of these species may have a naturally lower habitat affinity for forests. For example, the rodent *T. swinderianus*, recorded only in cashew orchards, is mostly found in open-area land cover types such as savannas and grasslands, but also in croplands (Child, 2016). Similarly, the semi-aquatic water mongoose *A. paludinosus*, more commonly recorded in cashew orchards, is a highly versatile species (Do Linh San *et al*., 2015), while the serval *Leptailurus serval*, recorded only in cashew orchards, is known to be a generalist carnivore associated with well-watered long-grass savanna as reported for southern African environments (Thiel, 2011).

Our analysis at the trophic guild level further revealed that herbivores and insectivores comprised the most sensitive guilds to this form of land-use change. Despite representing 20% of the mammal species (i.e., 5 species), herbivores are restricted to only ∼6% of the total number of mammal records, comprising the least frequently recorded trophic guild. Herbivore abundance was generally variable in forests but consistently low in cashew orchards, with herbivores absent from five out of the eight cashew orchards surveyed. This is likely due to the lower plant biomass in the cashew, which is further aggravated by the annual clearing made between January and May, aiming to facilitate the collection of the cashew nuts from the ground (Sousa et al., 2015). Similarly, herbivores were also the only mammal guild being negatively affected by the conversion of forests into cashew orchards in northern Guinea-Bissau (Rossinyol-Fernàndez *et al*., 2024). There, herbivore abundance in cashew orchards increased during the late rainy period, probably due to the increase in the undergrowth following rainfall (Bonser & Reader, 1995). The abundance of insectivores also declined from forest to cashew land cover types. Despite the low-intensity management of cashew orchards, it is possible that the lower structural complexity of cashew orchards resulted in a lower resource availability for insectivores (Stone *et al*., 2018). However, no differences in temperature or invertebrate abundance were observed between forests and cashew orchards in India (Komanduri *et al*., 2023).

On the contrary, the overall carnivore guild appeared to benefit from the presence of cashew orchards, where their abundance is higher. Although trophic specialisation is a key determinant of species’ resilience to land-use change (Newbold *et al*., 2019), not all carnivore species are resource specialists (Di Minin *et al*., 2016, Gorman *et al*., 2024). The carnivore guild includes a variety of species that range in their resource specificity (Heim *et al*., 2019). In this case, we are not able to infer that all carnivore species are being favoured by forest conversion into cashew orchards, but perhaps the most abundant species which are more likely to be those driving the pattern, namely the pardine genet *Genetta pardina* that accounted for 56.7% of the carnivore records and *A. paludinosus* that accounted for 28%, in contrast to the 15.3% records from the remaining carnivore species (i.e., *L. serval*, the common slender mongoose *Herpestes sanguineus*, *H. ichneumon*, *M. capensis* and the common genet *Genetta genetta*). Indeed, the presence of *G. pardina* has been reported in a wide variety of habitats, ranging from primary forests to suburban areas (Gaubert and Do Linh San, 2016). Similarly, despite being mainly restricted to riparian habitats and their vicinities, *A. paludinosus* has been commonly found in both native and non-native land cover types (Do Linh San et al., 2019). Furthermore, it is also possible that vegetation structure may be also playing a role, affecting not only prey availability but also predator hunting facility, which often determines carnivore abundance (Blaum *et al*., 2007). In this respect, carnivores may benefit from the simplified cashew orchards (Ray *et al*., 2005). Moreover, omnivore abundance remained unaffected by any of the variables considered in this study. This can be attributed to the broader resource utilization of omnivores compared to the other feeding guilds (Newbold *et al*., 2019). As for the carnivores, it is likely that *C. gambianus* and the striped ground squirrel *Xerus erythropus*, accounting for 63.1% and 17.7% of the omnivore species records, respectively, are driving these patterns. These findings are consistent with Rege & Lee (2023) and Vasconcelos *et al*. (2015), who also found that cashew orchards supported more generalist species than forests.

We expected that mammals would be negatively affected by increasing distance from forest edges. However, the distances between cashew sampling sites and the nearest forest edge (averaging 544 ± 443 m) did not appear to restrict their use by mammals. This may be partly due to the relatively short distances involved, which are well within the mobility range of most mammals in the study area (Gorman *et al*., 2024). Similarly, the apparent lack of a clear response to human activity may result from the comparable levels of human presence recorded in both forests and cashew orchards, which showed little variability (Table 1). This consistency of human activity across land cover types may have minimised its impact as a determining factor for mammal presence.

While this study represents an important first step in understanding how biodiversity is responding to the ongoing land-use change in West Africa, the sampling effort may have limited the robustness of our conclusions. The geographic area sampled within the CNP was relatively small, and the short distance between sampling sites (despite no spatial autocorrelation has been observed), combined with a reduced number of trapping days, further restricted the dataset. Consequently, the low sampling effort resulted in a low number of records per site particularly for the less recorded species, preventing a more comprehensive assessment of mammal responses to land-use changes. For instance, this prevented us from undertaking a species-specific approach. We recognise that by grouping mammal species in trophic guilds we are grouping species that might be ecologically different in aspects other than their diets (e.g., locomotion habit, diel activity, body size, habitat preferences, etc; Ferreira et al., 2018) and our guild-level approach might be influenced by the most recorded species, as discussed above in the case of the carnivore and omnivore guilds. In addition, although the number of camera-trap records is commonly used in mammal studies as a proxy of species abundance (Rossinyol-Fernàndez *et al*., 2024 and Goebel *et al*., 2025), this metric might be inflated by differential species detectability (e.g., species with smaller home ranges might be detected more often). Despite these limitations, our findings align with expectations, suggesting that a greater sampling effort would likely capture additional species in the forest land-cover type. This would further strengthen the present conclusions and underscore the significant impacts of cashew expansion on biodiversity in West Africa. Expanding the geographic scope and increasing the duration and intensity of sampling efforts could provide a more complete understanding of these dynamics.

### Conservation implications

Due to the high diversity within mammalian groups, responses to land-use change are often complex, context-dependent, and not uniformly negative. Forest-dependent species are more likely to face local extinction in disturbed land cover types (Palmeirim *et al*., 2020). Conversely, generalist and anthropophilic species may thrive in such environments, benefiting from increased resource availability and reduced predation pressure, as larger, less tolerant animals decline (Laurance & Vasconcelos, 2004; Niang *et al*., 2022). Examining changes in species assemblages while accounting for species-specific traits provides crucial insights into the broader impacts of land-use changes and ecosystem disturbances (Avenant & Cavallini, 2007). Promoting structural similarity between cashew orchards and adjacent forests may help sustain a greater diversity of mammals in these agricultural landscapes. For example, incorporating native trees into cashew orchards in India was shown to enhance bird diversity, underscoring the potential benefits of integrating elements of native vegetation into the monocultures (Madhu *et al*., 2024).

Our findings are in line with the general expectation that species assemblages undergo significant restructuring following land-use changes (Barnes *et al*., 2014, 2017). Such shifts can have detrimental effects on ecosystem functions and the services they provide (Lacher *et al*., 2019; Nhassé *et al*., 2024). For instance, herbivores contribute to nutrient cycling through grazing, while insectivores play a vital role in controlling insect populations. A decline in these trophic guilds could lead to the loss of critical regulatory functions and ecosystem services), ultimately impairing ecosystem resilience, functionality, and human well-being (Barnes *et al*., 2014). To promote mammal persistence, conservation strategies should aim to minimise forest conversion into cashew monoculture, while maintaining forest in adjacent areas, and preserving elements of the native vegetation within the plantations. This study provides novel insights into the impacts of land-use change on mammalian assemblages and offers valuable guidance for the development of conservation approaches in tropical regions facing rapid environmental transformation.

## Acknowledgements

We thank: the Institute of Biodiversity and Protected Areas (IBAP) of Guinea-Bissau for authorising and supporting the surveys, namely its director Aissa Regalla and CNP director Queba Quecuta; everyone who supported and welcomed us at the accommodation in Iemberém, namely Mamadú Baldé, Joia Tamba, Julmira Na Rescré, Hoina Na Cul, Mamadu Galisa, Djebu Seide, Abubácar Serra, Bubácar Na N’Caba, Cadidjatu Galisa, Aissatu Camará, Aissatu Turé, and Braima Camará, and all the landowners who allowed us to survey in their cashew farms; the CIBIO-CTM lab staff, namely Susana Lopes and Patrícia Ribeiro, for their support in the laboratory; and Mamadu Cassamá for his invaluable guidance in the field. AFP was supported by the European Union’s Horizon 2020 research and innovation programme under grant agreement No 854248 (TROPIBIO). This programme further funded the fieldwork.

## Author contributions

AFP conceived the ideas which were improved by comments from all authors; DN, JS, RO, IN, and AFP collected the data; DN and LP processed the camera-trap records; DN analysed the data and led the writing; and all authors contributed with comments and revisions to drafts of the manuscript.

## Data Availability

The data that support the findings of this study will soon be made available in a data paper. In the meantime, data will be made available on request.

