## Supplementary Material for "Mammal responses to habitat degradation induced by cashew expansion in West Africa"

**Table S1.** Sampling effort per sampling site used to survey mammal assemblages in the Cantanhez National Park, Guinea-Bissau. Site name, geographic coordinates, habitat type, name of the nearest village and number of trap-days (sampling effort) are indicated for each sampling site.

| | Geographic coordinates | | | | |
|---|---|---|---|---|---|
| **Site name** | **North** | **West** | **Habitat type** | **Nearest village** | **No. trap-days** |
| Cam1 | 11.20968 | 15.03850 | Forest | Cambeque | 30 |
| Cam2 | 11.20470 | 15.03853 | Forest | Cambeque | 30 |
| Cam3 | 11.19866 | 15.03882 | Forest | Cambeque | 30 |
| Cam4 | 11.19398 | 15.03956 | Forest | Cambeque | 30 |
| Lau1 | 11.21769 | 15.01977 | Forest | Lauchande | 32 |
| Lau2 | 11.21359 | 15.01813 | Forest | Lauchande | 32 |
| Lau3 | 11.20964 | 15.01949 | Forest | Lauchande | 32 |
| Lau4 | 11.20558 | 15.02135 | Forest | Lauchande | 32 |
| Mad1 | 11.24302 | 15.05483 | Forest | Madina | 30 |
| Mad2 | 11.24527 | 15.05775 | Forest | Madina | 30 |
| Mad3 | 11.25058 | 15.06073 | Forest | Madina | 29 |
| Mad4 | 11.25405 | 15.06513 | Forest | Madina | 30 |
| Cashew1 | 11.25760 | 15.07673 | Cashew orchard | Lauchande | 32 |
| Cashew2 | 11.25923 | 15.08227 | Cashew orchard | Lauchande | 32 |
| Cashew3 | 11.24141 | 15.05224 | Cashew orchard | Lauchande | 24 |
| Cashew4 | 11.24167 | 15.04611 | Cashew orchard | Lauchande | 32 |
| Cashew5 | 11.22479 | 15.01646 | Cashew orchard | Cambeque | 28 |
| Cashew6 | 11.22775 | 15.01821 | Cashew orchard | Cambeque | 30 |
| Cashew7 | 11.22791 | 15.02426 | Cashew orchard | Cambeque | 15 |
| Cashew8 | 11.23022 | 15.02710 | Cashew orchard | Cambeque | 30 |
| Cashew9 | 11.22206 | 15.03895 | Cashew orchard | Madina | 30 |
| Cashew10 | 11.21507 | 15.03737 | Cashew orchard | Madina | 30 |
| Cashew11 | 11.21808 | 15.04491 | Cashew orchard | Madina | 30 |
| Cashew12 | 11.22261 | 15.03015 | Cashew orchard | Madina | 29 |

**Table S2.** Mammal species list recorded in forests and cashew orchards across 24 sampling sites in the Cantanhez National Park, Guinea-Bissau. For each species we indicated the order, family, scientific name, common name, corresponding trophic guild (i.e., carnivores (CA), insectivores (IN), omnivores (OM) and herbivores (HE) and number (and corresponding percentage) of records in forests and cashew orchards.

| ORDER /Family | Scientific name | Common name | Trophic guild | Forests (%) | Cashew (%) |
|---|---|---|---|---|---|
| ARTIODACTYLA | | | | | |
| | Bovidae | | | | |
| | *Cephalophus silvicultor* | Yellow-backed duiker | HE | 1 (0.12%) | 0 |
| | *Cephalophus rufilatus* | Red-flanked duiker | HE | 9 (1.07%) | 0 |
| | *Philantomba maxwelli* | Maxwell's duiker | HE | 23 (2.73%) | 3 (0.36%) |
| | *Tragelaphus scriptus* | Bushbuck | HE | 8 (0.95%) | 4 (0.48%) |
| | Suidae | | | | |
| | *Potamochoerus porcus* | Red river hog | OM | 1 (0.12%) | 0 |
| CARNIVORA | | | | | |
| | Felidae | | | | |
| | *Leptailurus serval* | Serval | CA | 0 | 4 (0.48%) |
| | Herpestidae | | | | |
| | *Atilax paludinosus* | Marsh mongoose | CA | 6 (0.71%) | 44 (5.23%) |
| | *Herpestes ichneumon* | Egyptian mongoose | CA | 1 (0.12%) | 1 (0.12%) |
| | *Herpestes sanguineus* | Common slender mongoose | CA | 9 (1.07%) | 3 (0.36%) |
| | *Ichneumia albicauda* | White-tailed mongoose | IN | 146 (17.34%) | 50 (5.94%) |
| | *Mungos mungo* | Banded mongoose | IN | 0 | 1 (0.12%) |
| | *Mungos gambianus* | Gambian mongoose | IN | 1 (0.12%) | 3 (0.36%) |
| | Mustelidae | | | | |
| | *Mellivora capensis* | Honey badger | CA | 0 | 2 (0.24%) |
| | Viverridae | | | | |
| | *Civettictis civetta* | African civet | OM | 17 (2.02%) | 10 (1.19%) |
| | *Genetta pardina* | Pardine genet | CA | 36 (4.28%) | 62 (7.36%) |
| | *Genetta genetta* | Common genet | CA | 0 | 3 (0.36%) |
| PRIMATA | | | | | |
| | Cercopithecidae | | | | |

| | | | | |
|---|---|---|---|---|
| *Chlorocebus sabaeus* | Green monkey | OM | 8 (0.95%) | 4 (0.48%) |
| *Cercopithecus campbelli* | Campbell's monkey | OM | 7 (0.83%) | 0 |
| Hominidae | | | | |
| *Pan troglodytes verus* | Western chimpanzee | OM | 9 (1.07%) | 6 (0.71%) |
| RODENTIA | | | | |
| Sciuridae | | | | |
| *Funisciurus pyrropus* | Fire-footed rope squirrel | OM | 19 (2.26%) | 0 |
| *Heliosciurus gambianus* | Gambian sun squirrel | OM | 1 (0.12%) | 1 (0.12%) |
| *Xerus erythropus* | Striped ground squirrel | OM | 25 (2.97%) | 45 (5.34%) |
| Hystricidae | | | | |
| *Atherurus africanus* | African brush-tailed porcupine | IN | 11 (1.31%) | 7 (0.83%) |
| Nesomyidae | | | | |
| *Cricetomys gambianus* | Giant pouched-rat | OM | 173 (20.55%) | 77 (9.14%) |
| Thryonomyidae | | | | |
| *Thryonomys swinderianus* | Marsh cane-rat | HE | 0 | 1 (0.12%) |

**Table S3.** Results of the PERMANOVA and PERMDIST.

| **PERMANOVA** | **Df** | **Sum of Sqs** | **$R^2$** | **F** | **Pr(>F)** | |
|---|---|---|---|---|---|---|
| Group vector | 1 | 0.398 | 0.088 | 1.931 | 0.049 | |
| Residual | 20 | 4.119 | 0.912 | | | |
| Total | 21 | 4.517 | 1.000 | | | |
| **PERMIDST** | **Df** | **Sum Sq** | **Mean Sq** | **F** | **N.Perm** | **Pr(>F)** |
| Groups | 1 | 0.0001 | 0.0002 | 0.021 | 999 | 0.896 |
| Residuals | 20 | 0.164 | 0.008 | | | |

**Table S4.** Candidate Generalised Linear Models (GLMs) explaining mammal estimated species richness, overall species activity and activity of each trophic guild—carnivores, insectivores, omnivores, and herbivores—according to habitat structure ("Habitat"; given by the Principal Component 1 of the PCA including the variables: floor and understorey obstruction, density of lianas, palms and trees, tree species richness and height), landscape ("Dist"; Euclidian distance to the nearest forest) and degree of human activity ("Human"; number of camera-trap records on humans) across 24 sampling sites in the Cantanhez National Park, Guinea-Bissau. We provide the full set of alternative models, i.e., $\Delta AICc < 2$, ordered by AICc (Akaike Information Criterion for small samples) values. df = degrees of freedom; logLik = log-likelihood; $\Delta AICc = AICc_i - AICc_{min}$, $i = i^{th}$ model; $w_i$ = Akaike weights. As an exception, Generalised Additive Models have been used in the case of the activity of carnivores, in which we used the geographic coordinates ("Geogr.") as a smooth parameter.

| **Response** | **Intercept** | **Human** | **Dist** | **Habitat** | **offset** | **df** | **logLik** | **AICc** | **ΔAICc** | $w_i$ |
|---|---|---|---|---|---|---|---|---|---|---|
| *Observed species richness* | | | | | | | | | | |
| | 2.033 | | | | | 2 | -51.179 | 107.0 | 0 | 0.416 |
| | 2.029 | -0.097 | | | | 3 | -50.459 | 108.3 | 1.26 | 0.221 |
| | 2.033 | | | -0.028 | | 3 | -51.118 | 109.6 | 2.58 | 0.115 |
| | 2.033 | | -0.025 | | | 3 | -51.130 | 109.6 | 2.6 | 0.113 |
| | 2.028 | -0.095 | | -0.021 | | 4 | -50.425 | 111.2 | 4.21 | 0.051 |
| | 2.029 | -0.096 | -0.001 | | | 4 | -50.459 | 111.3 | 4.28 | 0.049 |
| | 2.033 | | -0.013 | -0.020 | | 4 | -51.110 | 112.6 | 5.58 | 0.026 |
| | 2.028 | -0.099 | 0.019 | -0.032 | | 5 | -50.408 | 114.6 | 7.58 | 0.009 |
| *Estimated species richness* | | | | | | | | | | |
| | 2.442 | | | -0.205 | | 3 | -63.683 | 134.7 | 0 | 0.272 |
| | 2.432 | -0.149 | | -0.191 | | 4 | -62.332 | 135.0 | 0.32 | 0.232 |
| | 2.448 | | -0.174 | | | 3 | -64.583 | 136.5 | 1.80 | 0.11 |
| | 2.449 | -0.174 | | | | 3 | -64.714 | 136.8 | 2.06 | 0.097 |
| | 2.462 | | | | | 2 | -66.197 | 137.0 | 2.33 | 0.085 |
| | 2.44 | | -0.072 | -0.163 | | 4 | -63.480 | 137.3 | 2.61 | 0.074 |
| | 2.441 | -0.134 | -0.141 | | | 4 | -63.650 | 137.7 | 2.95 | 0.062 |
| | 2.432 | -0.143 | -0.029 | -0.175 | | 5 | -62.297 | 138.3 | 3.65 | 0.044 |
| | -1.053 | | | | + | 2 | -68.267 | 141.2 | 6.47 | 0.011 |
| | -1.065 | | -0.112 | | + | 3 | -67.774 | 142.9 | 8.18 | 0.005 |
| | -1.061 | | | -0.064 | + | 3 | -68.100 | 143.5 | 8.83 | 0.003 |
| | -1.059 | -0.055 | | | + | 3 | -68.161 | 143.7 | 8.96 | 0.003 |
| | -1.067 | -0.028 | -0.105 | | + | 4 | -67.747 | 145.8 | 11.15 | 0.001 |
| | -1.065 | | -0.112 | <0.001 | + | 4 | -67.774 | 145.9 | 11.20 | 0.001 |

| | | | | | | | | | |
|---|---|---|---|---|---|---|---|---|---|
| -1.066 | -0.050 | | -0.061 | + | 4 | -68.012 | 146.4 | 11.68 | 0.001 |
| -1.067 | -0.028 | -0.104 | -0.003 | + | 5 | -67.747 | 149.2 | 14.55 | <0.001 |
| *Overall species abundance* | | | | | | | | | |
| 0.526 | | | | | 2 | -72.274 | 149.2 | 0 | 0.451 |
| 2.521 | | | -0.093 | | 3 | -71.983 | 151.3 | 2.12 | 0.156 |
| 2.522 | -0.098 | | | | 3 | -72.023 | 151.4 | 2.20 | 0.15 |
| 2.525 | | -0.039 | | | 3 | -72.221 | 151.8 | 2.60 | 0.123 |
| 2.519 | -0.087 | | -0.084 | | 4 | -71.785 | 153.9 | 4.74 | 0.042 |
| 2.521 | | 0.024 | -0.108 | | 4 | -71.970 | 154.3 | 5.11 | 0.035 |
| 2.522 | -0.093 | -0.018 | | | 4 | -72.012 | 154.4 | 5.20 | 0.034 |
| 2.518 | -0.098 | 0.050 | -0.113 | | 5 | -71.732 | 157.2 | 8.03 | 0.008 |
| *Carnivore abundance* | | | | | | | | | |
| 0.798 | | | 0.590 | | 3 | -42.396 | 92.1 | 0 | 0.545 |
| 0.793 | | 0.111 | 0.521 | | 4 | -42.233 | 94.8 | 2.69 | 0.142 |
| 0.790 | -0.115 | | 0.606 | | 4 | -42.245 | 94.8 | 2.72 | 0.14 |
| 0.888 | | 0.350 | | | 3 | -44.568 | 96.5 | 4.34 | 0.062 |
| 0.952 | | | | | 2 | -46.199 | 97.0 | 4.90 | 0.047 |
| 0.780 | -0.161 | 0.153 | 0.518 | | 5 | -41.952 | 97.7 | 5.53 | 0.034 |
| 0.876 | -0.176 | 0.393 | | | 4 | -44.278 | 98.9 | 6.78 | 0.018 |
| 0.952 | -0.038 | | | | 3 | -46.186 | 99.7 | 7.58 | 0.012 |
| *Insectivore abundance* | | | | | | | | | |
| 0.998 | | | -0.575 | | 3 | -47.571 | 102.5 | 0 | 0.391 |
| 0.932 | | 0.418 | -0.857 | | 4 | -46.197 | 102.7 | 0.27 | 0.342 |
| 0.989 | 0.160 | | -0.596 | | 4 | -47.327 | 105.0 | 2.53 | 0.110 |
| 0.930 | 0.077 | 0.399 | -0.853 | | 5 | -46.132 | 106.0 | 3.54 | 0.067 |
| 1.172 | | | | | 2 | -50.877 | 106.4 | 3.91 | 0.055 |
| 1.164 | | -0.111 | | | 3 | -50.759 | 108.9 | 6.38 | 0.016 |
| 1.169 | 0.077 | | | | 3 | -50.832 | 109.0 | 6.52 | 0.015 |
| 1.159 | 0.110 | -0.132 | | | 4 | -50.673 | 111.7 | 9.22 | 0.004 |
| *Omnivore abundance* | | | | | | | | | |
| 1.792 | | | | | 2 | -59.521 | 123.7 | 0 | 0.366 |
| 1.773 | -0.227 | | | | 3 | -58.642 | 124.6 | 0.94 | 0.228 |
| 1.782 | | -0.147 | | | 3 | -59.080 | 125.5 | 1.82 | 0.147 |
| 1.792 | | | -0.022 | | 3 | -59.511 | 126.4 | 2.68 | 0.096 |
| 1.769 | -0.197 | -0.098 | | | 4 | -58.449 | 127.3 | 3.58 | 0.061 |
| 1.773 | -0.228 | | 0.009 | | 4 | -58.641 | 127.6 | 3.96 | 0.050 |
| 1.778 | | -0.215 | 0.112 | | 4 | -58.920 | 128.2 | 4.52 | 0.038 |
| 1.766 | -0.195 | -0.165 | 0.107 | | 5 | -58.298 | 130.3 | 6.67 | 0.013 |
| *Herbivore abundance* | | | | | | | | | |
| 0.794 | | | -0.774 | | 3 | -20.747 | 48.8 | 0 | 0.452 |
| -0.526 | | | | | 2 | -23.089 | 50.8 | 1.98 | 0.168 |
| -0.803 | -0.093 | | -0.785 | | 4 | -20.711 | 51.8 | 2.95 | 0.103 |
| -0.793 | | 0.025 | -0.788 | | 4 | -20.745 | 51.8 | 3.02 | 0.100 |

| | | | | | | | | | |
|---|---|---|---|---|---|---|---|---|---|
| | -0.643 | | -0.525 | | 3 | -22.329 | 52.0 | 3.16 | 0.093 |
| | -0.528 | -0.070 | | | 3 | -23.074 | 53.5 | 4.65 | 0.044 |
| | -0.644 | 0.046 | -0.537 | | 4 | -22.321 | 55.0 | 6.17 | 0.021 |
| | -0.803 | -0.106 | 0.064 | -0.823 | 5 | -20.704 | 55.2 | 6.33 | 0.019 |

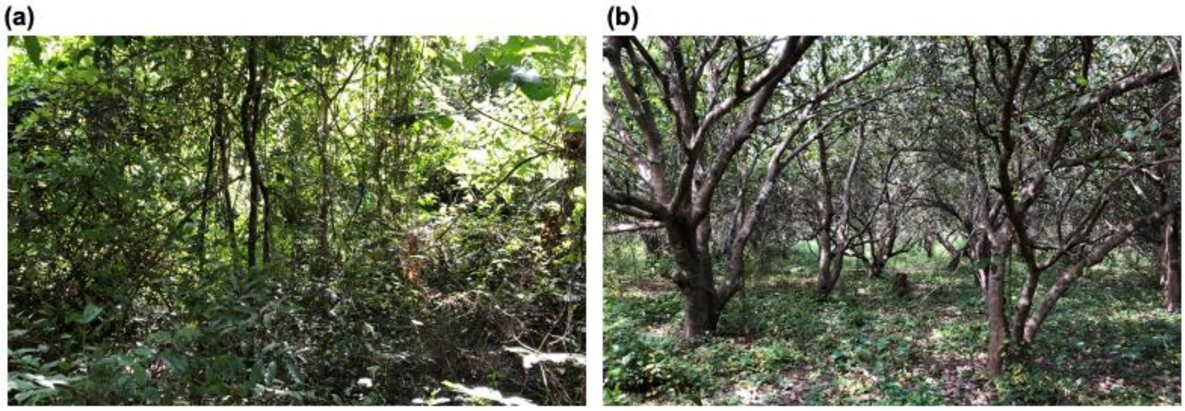


**Figure S1.** Photographs illustrating the two land-use types surveyed in the Cantanhez National Park, Guinea-Bissau: (a) closed-canopy forests and (b) cashew orchards.

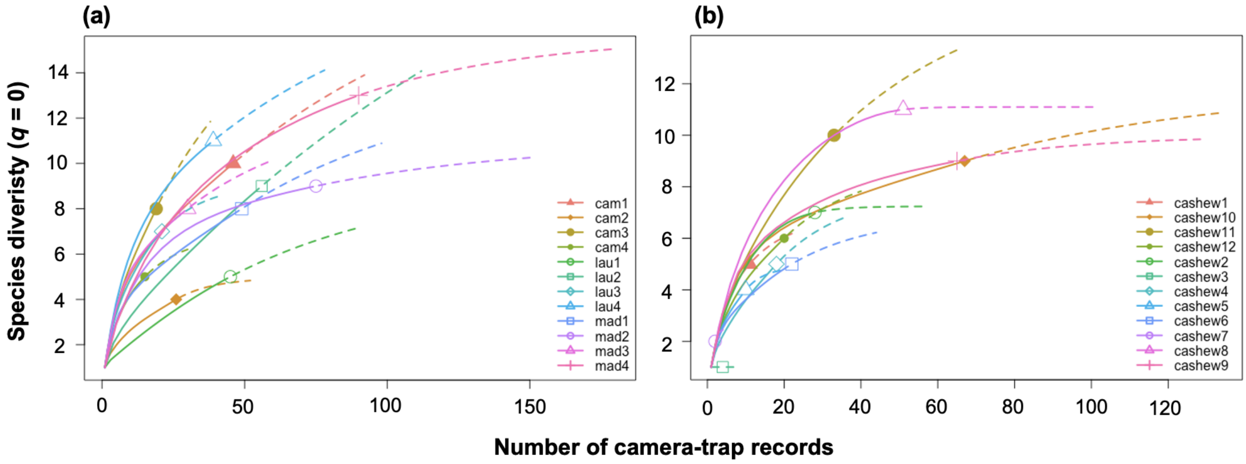


**Figure S2.** Individual-based extrapolation curves showing mammal species richness (q = 0) surveyed across 24 sampling sites in the Cantanhez National Park, Guinea-Bissau. Solid lines represent rarefaction, and dashed lines represent extrapolation. The solid dots, triangles and squares represent the reference samples, i.e., the cumulative number of mammal records at each sampling site. Each curve was extrapolated to double of the number of individuals recorded in the corresponding sampling site.

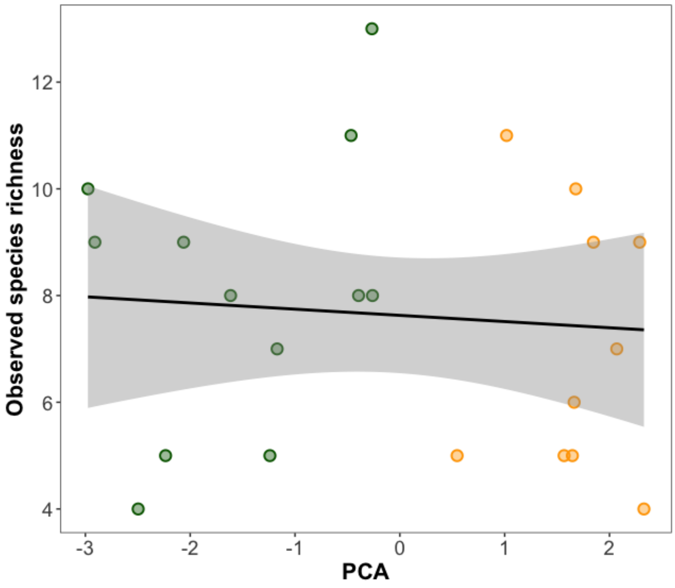


**Figure S3.** Relationship between observed species richness and the local habitat characteristics as denoted by the scores of the first component of the Principal Component Analysis (CP1). The shaded area represents the 95% confidence region; solid dots indicate observed values. The line represents the model adjusted. Dots are colour coded according to the land-cover type where the sampling sites were located (forests in green and cashew orchards in orange).
